# Dual cholinergic mechanisms for sculpting striatal dopamine in vivo

**DOI:** 10.64898/2025.12.19.695021

**Authors:** Dylan R. Flink, Nicholas G. Faturos, Haowei Zhang, Arif A. Hamid

## Abstract

Striatal dopamine (DA) and acetylcholine constitute a computationally powerful neuromodulatory dyad that orchestrates action selection, motivational vigor, and reward learning. Striatal cholinergic interneurons (CINs) synapse onto DA axons and stimulate DA release via β_2_-containing nicotinic receptors (β_2_-nAChRs), providing a local modulatory channel distinct from midbrain-derived spikes. Yet whether this mechanism operates *in vivo* and how it contributes to striatal computations, orthogonal to those facilitated by prediction errors encoded in the DA somata, remains unresolved. Using spatiotemporally precise optogenetic stimulation of CINs paired with ultrafast DA imaging in the dorsal striatum, we identify a biphasic mode of CIN-evoked DA release in awake mice. CIN activation evoked an ultra-fas β_2_-nAChR-dependent DA transient peaking within ~40 ms that requires millimeter-scale CIN synchrony and is systematically obscured by conventional imaging bandwidths or slow DA sensor kinetics. A second, delayed DA elevation peaking at ~180 ms persisted despite β_2_-nAChR blockade and is mediated by muscarinic receptors, α_7_- and α_6_*-nAChRs, and a non-cholinergic CIN messenger, revealing a previously unrecognized polysynaptic CIN mechanism. Our observations reconcile discrepancies between slice and *in vivo* studies and show that CINs engage DA release through two mechanistically distinct pathways: (i) an ultra-fast, monosynaptic mode provides a spatially expansive DA signal ideally positioned to reconfigure striatal microcircuits globally, and (ii) a slower, spatially flexible polysynaptic DA mode suited for sustaining ongoing decisions. This dual architecture expands the computational repertoire of striatal DA dynamics in behavioral and cognitive control.

## Introduction

Circuit mechanisms that coordinate dopamine (DA) release across brain regions and timescales are essential for flexible behavioral control and present as vulnerable substrates for neurological and psychiatric disorders^1–4^. Beyond the canonical view that DA release dynamics are shaped by midbrain somatic spikes, a series of *ex vivo* studies have revealed a parallel mode of DA regulation entirely within the striatum, mediated by local microcircuits^5,6^. In particular, cholinergic interneurons of the striatum (CINs) synapse onto DA axons^7^ and evoke axonal action potentials^8^ to drive DA release via β_2_-subunit containing nicotinic receptors (β_2_-nAChRs)^9,10^. This acetylcholine-evoked DA release persists in preparations lacking DA somata, is abolished by the β_2_-nAChR antagonist dihydro-β-erythroidine hydrobromide (DHβE), and leverages the same exocytotic machinery as soma-driven DA efflux^8^. This mechanism, therefore, positions CINs as spatiotemporally precise local modulatory hubs that gate cortical, thalamic, and striatal influences directly onto DA axons^9,11–15^, bypassing the midbrain as the sole command center for DA release.

This local mode of striatal DA control has important implications for reinforcement learning (RL) theory, motivation and cognitive control. Classical RL frameworks cast midbrain DA neurons as a unified source of reward prediction errors (RPEs)^16–18^, and posit that their broad axonal arbors^19,20^ obligate a globally uniform DA release that encodes this learning signal^21–24^. By contrast, emerging perspectives argue that striatal DA fluctuations multiplex RPEs with additional decision variables, of unknown circuit origin, that support ongoing motivational, choice, and cognitive control processes^25–33^. In this view, CINs are hypothesized to act as circuit-level inference nodes^34,35^ that could translate cortical conflict signals and local circuit entropy into a rapid, regional rerouting of DA release during the performance epoch^32,36,37^. The local cholinergic control of DA via β_2_-nAChRs, therefore, provides a concrete candidate mechanism for sculpting regional DA dynamics in service of performance and delay epoch circuit computations before rewards materialize^25–33^, orthogonal to midbrain-derived RPEs that predominantly guide value learning at the moment of reward delivery^16–18,25–33^. If empirically supported, such a circuit architecture would implement a modular RL algorithms mobilized under specific task epochs and behavioral demands by leveraging a collaborative exchange of striatal -vs- midbrain control over DA release in striatum^31^.

Despite strong *ex vivo* evidence and compelling computational motivation, whether CINs locally shape DA dynamics in awake, behaving animals is controversial^4,6,38^. Recent *in vivo* studies relied exclusively on loss-of-function approaches rather than direct CIN activation: genetic deletion of β_2_-nAChRs on DA axons or systemic DHβE administration failed to alter spontaneous or reward-evoked DA fluctuations^39,40^. These negative findings raise the possibility that unknown circuit interactions override monosynaptic cholinergic modulation of DA selectively *in vivo*, or that the consequences of CIN-DA control were obscured by technical limitations of these studies^37,38^. Consequently, the circuit logic by which striatal microcircuits exert real-time control over DA to regulate ongoing motivational and cognitive control is fundamentally unresolved.

## Results

### CIN activation triggers a biphasic DA release in vivo

We sought to clarify these inconsistencies by directly stimulating CINs and monitoring the consequent DA release in the striatum of awake head-restrained mice. ChAT-Cre animals were injected with the red-shifted ChrimsonR opsin (AAV5-syn-FLEX-ChrimsonR) and green DA sensor, dLight1.2 (AAV5-hsyn-dLight1.2) in the dorsal striatum (DS) (**Fig. 1a**). We implanted a 3-mm diameter cannula over the DS^36^ and imaged dLight during spatiotemporally precise CIN stimulation (**Fig. 1b**). This approach avoided the recently reported^37,38^ optical artifact caused by blue-light excitation of red DA sensors, while uniquely allowing synchronous activation of CINs across the entire DS.

**Figure 1:**
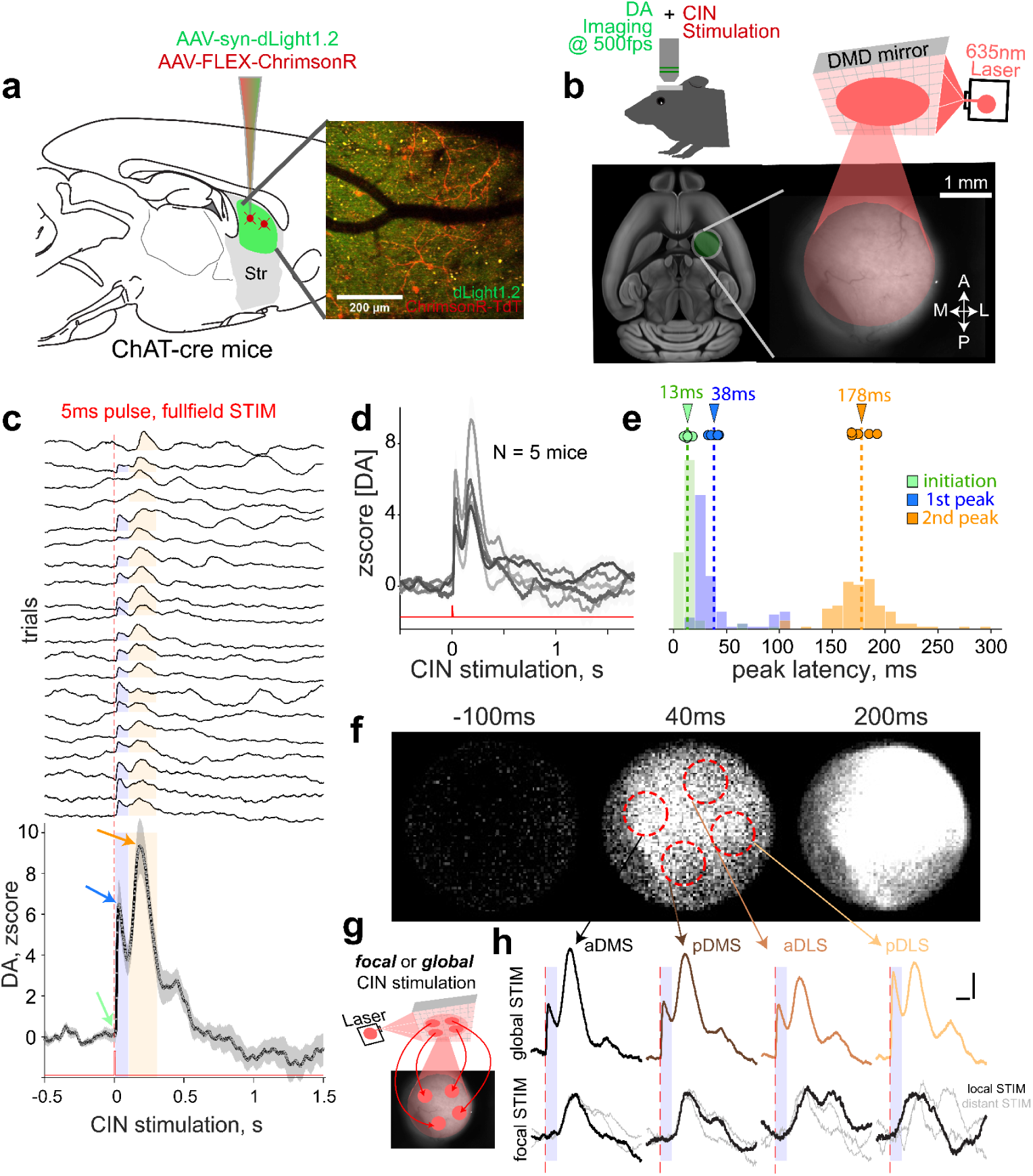
A biphasic DA release following global CIN stimulation In Vivo. **(a)** Schematic of methods used to achieve CIN-specific expression of ChrimsonR and broad dLight1.2 delivery into the dorsal striatum (DS). *Right:* An example of opsin and sensor expression in an awake mouse. **(b)** After cannula implantation, head-fixed animals were imaged at 500 fps while DS CINs were stimulated using a digital micromirror device (DMD) that projected a 635 nm laser either across the entire imaging field or onto restricted DS regions. *Bottom:* Example widefield field of view (FOV) aligned to the Allen CCF^41^. **(c)** A 5-ms laser pulse triggers a biphasic DA release. *Top*: trial-by-trial dLight fluorescence averaged across all DS pixels. *Bottom*: Session-averaged trace with circles indicating DA level at each acquisition frame. Data displayed as Mean ± SEM. Blue shading indicates the quantification window (0-0.1 s) for the “first peak”; yellow shading indicates the 0.1-0.3 s window for the “second peak”. **(d)** Mean traces across DS in five animals examined, demonstrating a biphasic response. **(e)** Quantification of average latency relative to laser onset, with colors matching the arrows in panel **c**. Green indicates the latency at which the signal rises significantly above baseline; blue and orange correspond to the latencies of the first and second peaks, respectively. Histograms show individual trials pooled across animals; circles above represent within-animal means. **(f)** Spatial spread of CIN-evoked DA during full-field DS stimulation in an example mouse. Single frames just before, at the first peak, and at the second peak (−100 ms, 40 ms, and 200 ms relative to laser onset) Dotted redlines indicate ROIs for both fluorescence quantification (in panel **h**) and location of laser spots (in panel **g**). **(g)** Five laser delivery strategies used to manipulate stimulation area: either global (full field, 7 mm^2^) or spot stimulation restricted to DS quadrants (0.5 mm^2^ each). **(h)** *Top row:* average DA responses detected at the anterior dorsomedial (aDMS), posterior dorsomedial (pDMS), anterior dorsolateral (aDLS), or posterior dorsolateral (pDLS) striatum during global laser activation. Note that both DA peaks are present at all locations. *Bottom row*: DA responses when smaller laser spots targeted the same region where fluorescence was measured (“local STIM,” dark traces) versus when the laser targeted a distant DS quadrant (“distant STIM,” gray traces). Note that smaller laser spots evoked only the delayed second DA elevation, including at non-targeted locations. Shaded blue boxes indicate the epoch where the first peak is typically (0 - 0.1s) detected relative to laser onset (dotted red line), as in panel **c**. Scale bars at the top right indicate 100ms horizontally and 2 standard units of DA elevation.

We first activated CINs with a single, 5-ms full-field red-light pulse (7.5 mW/mm^2^ across the 7mm^2^ FOV) and imaged dLight fluorescence at 500 frames per second. We observed a distinct biphasic DA response in DS in every animal examined (**Fig. 1c-d**). This DA elevation began within 13 ± 1.3 ms (Mean ± SEM, N = 5 mice) of laser onset, and an initial increase peaked at 38 ± 1.8 ms synchronously across DS (**Fig. 1e**). After a brief decline, a secondary DA elevation peaked at 178 ± 4.3 ms and was also evident across the DS, with little peak-time jitter (**Fig. 1e-f**). Surprisingly, delivering power-density and duration-matched laser pulses to smaller DS territories (0.5 mm^2^) elicited only the second DA elevation at the site of laser illumination that was also evident in distant, unstimulated DS regions (**Fig. 1g-h**). These observations establish that CIN activation *in vivo* evokes distinct modes of DA release: an ultra-fast, globally synchronous DA efflux and a delayed transient that spreads beyond its initiation site, suggesting dissociable underlying mechanisms support these DA responses.

### Ultra-fast DA peaks are selectively mediated by β_2_-nAChRs

Previous *ex vivo* studies have demonstrated that CIN activation can drive *monosynaptic* DA release via β_2_-nAChRs, with DA axons firing an action potential approximately 10 ms after CIN photoactivation^8^, and the resulting DA efflux peaking 20-30 ms later^42^. Given the similarly rapid onset and peak latency of the first CIN-evoked DA peak we observed *in vivo*, we hypothesized that this component reflects the same monosynaptic β_2_-nAChR-mediated mechanism described *ex vivo*^7–10,42^. By contrast, the markedly slower secondary DA peak suggested a distinct mechanism, potentially arising from rebound excitation of CINs or polysynaptic interactions that recruit additional cell types and cholinergic receptor classes. Accordingly, we predicted that inhibiting β_2_-nAChRs with DHβE would abolish the first CIN-evoked DA peak, and possibly the second if a rebounding CIN activation similarly relied on β_2_-nAChRs again.

Consistent with the first prediction, systemic DHβE (1 mg/kg, i.p.) selectively eliminated the first DA peak evoked by full-field CIN stimulation (**Fig. 2a-b**). This effect was also observed in the presence of the non-selective nAChR antagonist, mecamylamine (**Fig. 2c-d**), confirming that the initial, ultra-fast peak is a mono-synaptic CIN-evoked DA release mediated by β_2_-nAChRs (hereafter referred to as ^β_2_-nAChR^[DA]).

**Figure 2:**
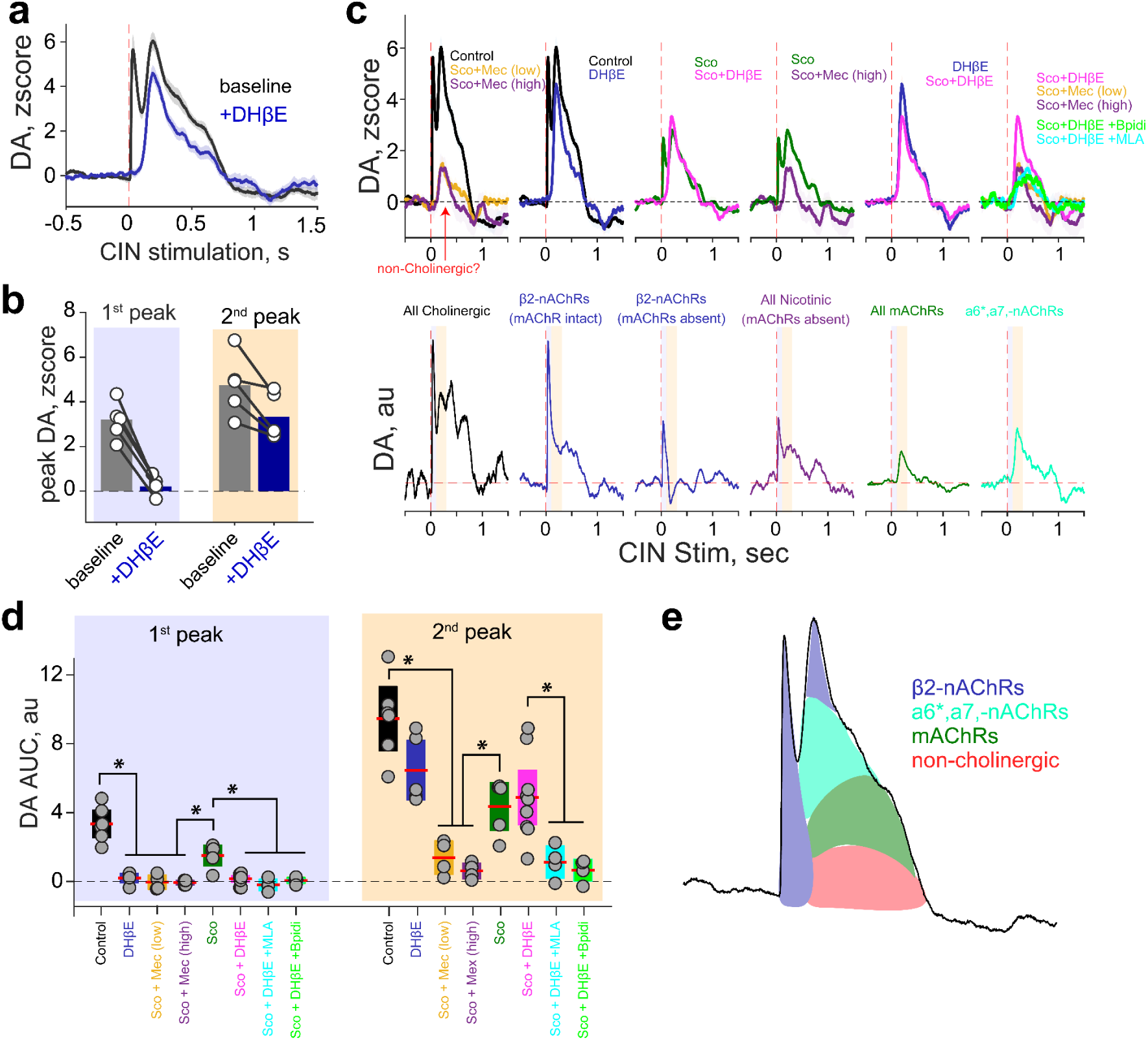
Pharmacological profiling reveals distinct mechanisms of CIN-evoked DA release. **(a)** Average DA across dorsal striatal FOV evoked by full-field CIN stimulation under baseline conditions and following systemic β_2_-nAChR antagonism (DHβE, 1 mg/kg, i.p.). These traces are averaged across all trials for each animal (N = 20 trials, 5 mice per condition). Data displayed as Mean ± SEM. **(b)** Quantification of peak evoked DA within time windows indexing the first (0 - 0.1 sec; blue) and second (0.1 - 0.3 sec; yellow) peaks illustrated as shaded regions in Fig. 1c. Data displayed as Mean ± SEM. DHβE selectively abolished the first CIN-evoked DA peak (two-way rmANOVA, significant main effect of DHβE: F(1,4) = 24.35, P = 0.002; drug x peakTime interaction: F(1,4) = 3.1, P = 0.01; post-hoc paired t-test: Early peak magnitude is different between control vs DHβE: t(8) = 7.326, P = 0.0001), while producing only a modest reduction in the second peak (post-hoc paired t-test: Late peak magnitude was not different between control vs DHβE: t(8) = 1.852, P = 0.10). **(c)** *Top:* Population-averaged CIN-evoked DA timecourse across all animals (N = 5 mice) under various systemic pharmacological manipulations. All data displayed as Mean ± SEM, aligned to laser onset. Each panel examines the respective contributions of various cholinergic receptor subtypes. *First panel*: contribution of all cholinergic receptors by systemic injection of a cocktail containing muscarinic antagonist scopolamine (Sco) and mecamylamine (Mec), either at low (1mg/kg each) or high (5mg/kg) doses. Note a non-cholinergic DA peak that is observed with the drug cocktail on board at either dosage. *Second panel*: contribution of β_2_-nAChRs with muscarinic receptors intact displayed in panel **a**. *Third panel*: contributions of β_2_-nAChRs without muscarinic receptors available by comparing timecourse of evoked DA between administration of scopolamine (Sco; 1mg/kg) or a cocktail of scopolamine and DHβE (1mg/kg each). *Fourth panel*: contributions of all nicotinic receptors examined by contrasting scopolamine (1mg/kg) alone against a cocktail of scopolamine and mecamylamine (1mg/kg each). *Fifth panel*: contributions of muscarinic receptors by comparing DHβE (1mg/kg) against a cocktail of scopolamine and DHβE (1mg/kg each). *Sixth panel*: contributions of non-DHβE sensitive nicotinic receptors by contrasting DA timecourse of scopolamine and DHβE (1mg/kg each) and scopolamine (Sco) and mecamylamine (Mec) at two doses. Directly testing the contributions of α_7_- or α_6_*-nAChRs with their respective antagonists (MLA or bPiDl, 1 mg/kg each) administered in a cocktail with scopolamine and DHβE (1mg/kg each) reduces evoked DA to a level indistinguishable from scopolamine (Sco) and mecamylamine (Mec). *Bottom panels* emphasize these contributions by directly subtracting the pharmacological conditions to isolate the contributions of each cholinergic receptor class. Shaded regions indicate windows where first and second peaks were detected, same as Fig. 1c. **(d)** Quantification of pharmacological effects on evoked DA release area under the curve (AUC) of the predefined time windows indexing the first (0 - 0.1 sec; blue) and second (0.1 - 0.3 sec; yellow) peaks illustrated as shaded regions in Fig. 1c. Each dot is average for each mouse tested, and the box plots show mean across animals (red horizontal line) and 95% confidence interval (top and bottom box boundaries). Pharmacological manipulations differentially affect the early versus late components of CIN-evoked DA release (two-way rmANOVA, significant main effect of Drug (F(7,14) = 19.65, P = 3.15 × 10^−6^), a significant main effect of PeakTime (F(1,2) = 69.53, P = 0.014), and a significant Drug × PeakTime interaction (F(7,14) = 12.04, P = 5.77 × 10^-5^). Post hoc paired t-test comparisons for each drug condition are indicated with a star for significance, shown below. Early peak AUC difference: Control vs Sco+Mec: t(3) = 8.083, P = 0.004; Control vs DHβE: t(4) = 10.332, P = 0.0004951; Sco vs Sco+DHβE: t(3) = 6.333, P = 0.00796; Sco vs Sco+Mec: t(3) = 3.948, P = 0.02899; DHβE vs Sco+DHβE: t(4) = 1.000, P = 0.3739; Sco+DHβE vs Sco+Mec: t(3) = 0.816, P = 0.4741; Sco+DHβE+MLA vs Sco+Mec: t(2) = 0.525, P = 0.6518; Sco+DHβE+bPiDl vs Sco+Mec: t(2) = 1.728, P = 0.2261. Late peak AUC difference: Control vs Sco + Mec: t(3) = 6.976, P = 0.006044; Control vs DHβE: t(4) = 3.671, P = 0.02138; Sco vs Sco+DHβE: t(3) = −1.071, P = 0.3627; Sco vs Sco+Mec: t(3) = 7.904, P = 0.004221; DHβE vs Sco+DHβE: t(4) = 1.000, P = 0.3739; Sco+DHβE vs Sco+Mec: t(3) = 3.283, P = 0.04633; Sco+DHβE+MLA vs Sco+Mec: t(2) = −0.056, P = 0.9607; Sco+DHβE+bPiDl vs Sco+Mec: t(2) = −3.043, P = 0.09316. **(e)** Schematic summarizing pharmacological experiments to isolate the contributions of various receptors to the biphasic cholinergic evoked DA release.

The second DA peak was mildly reduced by DHβE (**Fig. 2b**, *right*), but remained robustly detectable, indicating that it does not rely primarily on β_2_-nAChRs. In contrast, CIN-evoked DA release is completely abolished by DHβE in brain slices^8,9,43,44^, demonstrating that the secondary peak does not arise from the same monosynaptic axo-axonal ^β_2_-nAChR^[DA] mechanism. This persistence with DHβE also makes a rebound CIN activation that re-engages β_2_-nAChRs an unlikely origin of the second peak. Instead, this DA event is most consistent with *in vivo* recruitment of an intact polysynaptic striatal-midbrain circuit that is severed or functionally absent in the slice preparation.

To examine whether muscarinic acetylcholine receptors mediate this delayed DA response, we administered scopolamine in a cocktail with mecamylamine at low (1mg/kg each, i.p.) or high (5mg/kg each, i.p.) doses. Broadly antagonizing both cholinergic receptors precipitously reduced laser-evoked DA release (**Fig. 2c**, *left*), although a small, yet significant, delayed peak persisted at both dosages, pointing to a non-cholinergic CIN messenger^45–47^ that can evoke DA release *in vivo*.

Administering scopolamine alone reduced CIN-evoked DA, although both are still detected (**Fig. 2c**, *middle*), suggesting that muscarinic receptors potentiate ^β_2_-nAChR^[DA] and are not the sole mediators of the delayed DA peak. The latter notion is further supported by scopolamine and DHβE co-administration yielding a larger second DA peak than scopolamine/mecamylamine cocktail (**Fig. 2c**, *middle*), implicating the involvement of DHβE-insensitive nAChRs. In the striatum, α_7_- and α_6_*-nAChRs mediate CIN regulation of glutamatergic afferents from cortex and thalamus^48–51^, so we administered their antagonists (MLA 1mg/kg i.p and bPiDl 1mg/kg i.p, respectively) in a cocktail with DHβE and scopolamine to examine their contribution. We found that either drug reduced CIN-evoked DA release to a level indistinguishable from that produced by the scopolamine/mecamylamine cocktail (**Fig. 2c**, *right*), suggesting a synergistic contribution of α_7_- and α_6_*-nAChRs in promoting the second DA peak. Together, our pharmacological studies reveal that the delayed DA elevation is evoked by redundant cholinergic mechanisms involving β_2_-nAChRs, mAChRs, α_7_-nAChRs, α_6_*-nAChRs, and a non-cholinergic transmitter released by CINs (**Fig. 2d-e**). In contrast, the ultra-fast ^β_2_-nAChR^[DA] peak reflects a distinct, monosynaptic mode of cholinergic DA release *in vivo*.

### CIN-evoked ^β2-nAChR^[DA] is masked under standard imaging conditions

Previous studies^39,40^ that did not detect β_2_-nAChR regulation of DA dynamics relied on 10-20 Hz fiber photometry and fluorescent sensors with slow kinetics, likely obscuring the ultra-fast ^β_2_-nAChR^[DA] peak observed here. We reasoned that such bandwidth limitations would blur the distinct phases of CIN-evoked DA peaks into a unitary transient, thereby masking the contribution of β_2_-nAChRs. To test this directly, we imaged CIN-evoked DA using dLight1.2 at a reduced acquisition rate of 16 fps, matching the temporal bandwidth of conventional photometry. Under these conditions, full-field CIN stimulation still evoked a significant DA increase that was synchronous across the DS (**Fig. 3a-b**), but the biphasic dynamics evident at 500 fps were not resolved. Systemic DHβE reduced the magnitude and slowed the rising phase of this unitary response (**Fig. 3c-e**), but did not abolish evoked DA, indicating that temporal undersampling aliases the fast ^β_2_-nAChR^[DA] component into a unitary CIN-evoked elevation.

**Figure 3:**
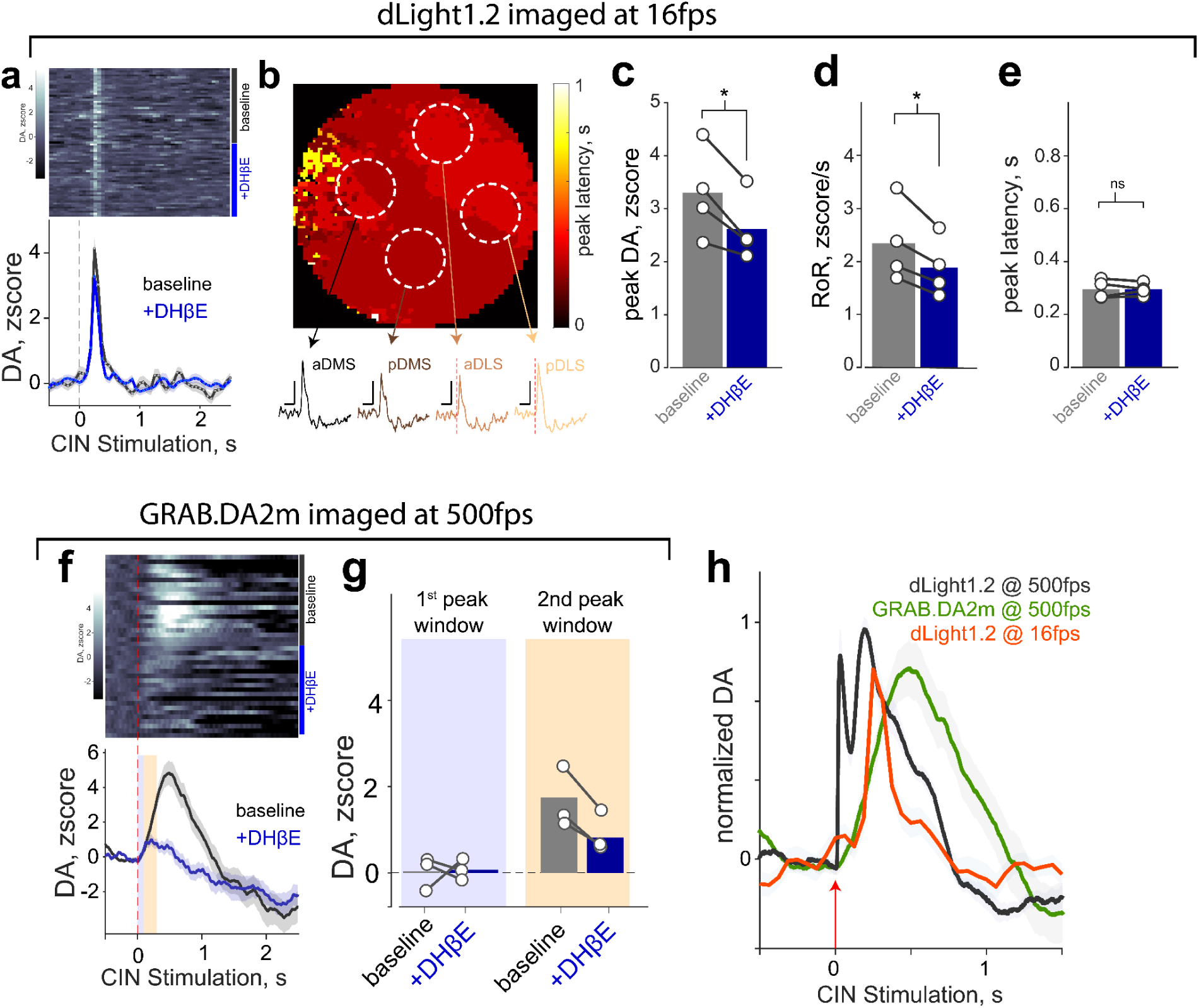
Slow sampling and sensor kinetics alias the ultra-fast β2-nAChR DA peak. **(a)** Heatmap (*top*) and mean trace (*bottom*) showing CIN-evoked DA release measured with dLight1.2 at 16 fps under before (“baseline”, black) and after systemic DHβE (1 mg/kg, blue), demonstrating that slow sampling reveals only a single DA elevation. **(b)** Spatial map of DA peak latencies across the dorsal striatum (DS) for an example mouse, with representative traces from striatal quadrants, illustrating synchronous DA recruitment. **(c)** DHβE significantly reduced the peak (one-way rmANOVA, significant main effect of DHβE: F(1,3) = 18.1, P = 0.02), **(d)** rate of rise (one-way rmANOVA, significant main effect of DHβE: F(1,3) = 19.88, P = 0.02), **(e)** without affecting the peak latency (one-way rmANOVA, significant main effect of DHβE: F(1,3) = 0.01, P = 0.89) of CIN-evoked DA release. **(f)** CIN-evoked DA quantified using GRAB.DA2m at 500 fps (heatmap and trace) reveals only a single sluggish DA elevation that peaks outside of the peak windows shown in Figure 1c (blue and orange shaded regions) for both baseline and DHβE conditions. Data displayed as Mean ± SEM. **(g)** Quantification of DHβE effects at the anticipated peak windows for the first (0 - 0.1s) and second peaks (0.1-0.3s), as in Fig. 1c. DHβE did not affect the peak magnitudes of GRAB.DA2m at the windows expected for first and second peaks (two-way rmANOVA, main effect of DHβE: F(1,2) = 3.63, P = 0.19; drug x peakTime interaction: F(1,2) = 3.87, P = 0.18. By contrast, when comparing the DA area under the curve (AUC) in a 0-1sec window, DHβE significantly attenuated CIN-evoked DA quantified by GRAB.DA2m (one-way rmANOVA, main effect of DHβE: F(1,2) = 23.8, P = 0.03). **(h)** Direct comparison of the DA timecourses when recorded with dLight1.2 at 500 fps (black), dLight1.2 at 16 fps (orange), and GRAB-DA2m at 500 fps (green). Note that slow frame rates or slow-kinetic sensors each collapse the biphasic DA response into a single broad transient.

We next asked whether utilizing fluorescent DA sensors with slower kinetics is sufficient to mask ^β_2_-nAChR^[DA], even under high-speed imaging. Under identical stimulation conditions, we imaged a separate cohort of mice expressing the D2R-based sensor GRAB-DA2m, which has a reported rise time of 40 ms, compared with 9.5 ms for the D1R-based dLight1.2^52–54^. CIN stimulation, while imaging GRAB-DA2m at 500 fps, produced a single sluggish DA transient peaking at 485 ± 3 ms, with no evidence of a fast component (**Fig. 3f-h**). DHβE significantly reduced the amplitude of this GRAB-DA2m response, but the remaining signal retained a monophasic profile (**Fig. 3f-g**). These findings demonstrate that slow sensor kinetics alone can mask the detection of ^β_2_-nAChR^[DA], even under high-speed imaging.

Taken together, our observations reveal that the ultra-fast β_2_-nAChR-dependent DA elevation can be systematically masked by restricted acquisition bandwidth and slow sensor kinetics. Either constraint alone is sufficient to alias the fast nicotinic component into a unitary CIN-evoked DA transient (**Fig. 3h**), preventing its identification as a distinct physiological event. We therefore conclude that previous attempts^39,40^ to identify β_2_-nAChR contributions to local DA modulation by CINs were constrained by technical limitations rather than the absence of this circuit mechanism *in vivo*.

### ^β_2_-nAChR^[DA] exhibits distinct temporal and spatial signatures

Having established β_2_-nAChR-dependent DA release *in vivo*, we next characterized its spatiotemporal features to facilitate their use as a signature for identifying ^β_2_-nAChR^[DA] events outside of optogenetic stimulation. We began by comparing the rise time and peak width of ^β_2_-nAChR^[DA] to the slower, non-β_2_-nAChR-dependent DA elevation across DS. In response to single pulse optogenetic stimulation, the ^β_2_-nAChR^[DA] component exhibited a markedly faster rising rate and narrower full width at half maximum (FWHM) (**Fig. 4a**). These DA kinetics parameters formed distinct clusters that were significantly separated (**Fig. 4b**). This separation indicates that ^β_2_-nAChR^[DA] occupies a distinct dynamical regime of striatal DA fluctuations, which would enable reliable classification.

**Figure 4:**
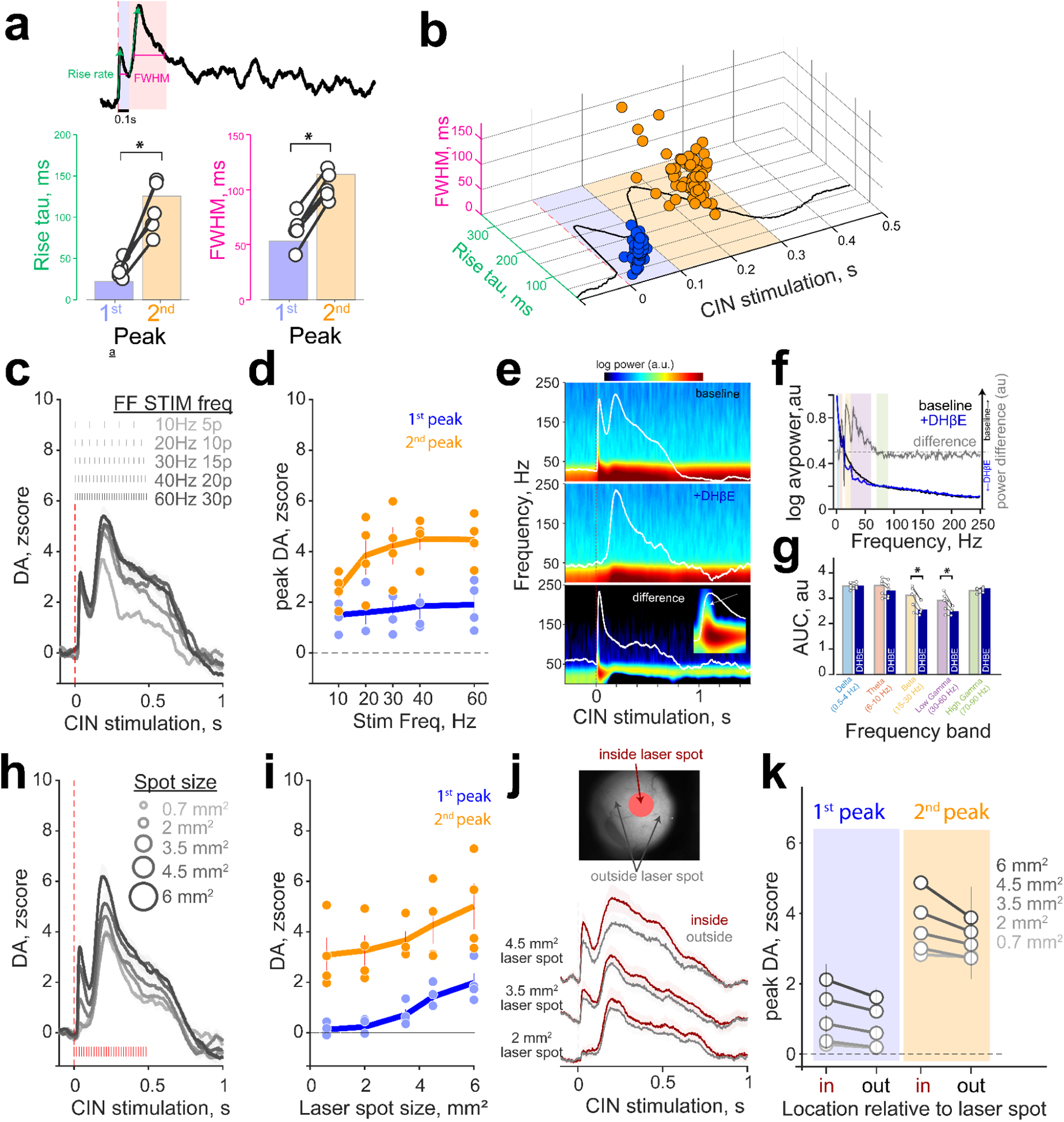
CIN-evoked DA peaks can be distinguished by separable spatiotemporal features. **(a)** *Top*: A representative single trial example of CIN-evoked DA illustrating the quantification of the rate of rise (green) and full width at half maximum (FWHM). *Bottom*: These kinetics measures are significantly different between the two cholinergic DA responses (N = 5 mice; one-way rmANOVA, main effect of peakTime on RiseTau: F(28.54), P =0.005; one-way rmANOVA, main effect of peakTime on RiseTau: F(115.51), P =0.0004). **(b)** Scatter plot showing the clustering of rise rate and width for the early (blue) and late (yellow) CIN-evoked DA peaks. Mahalanobis distance test for cluster separability between 2D distances of blue and yellow data: DM = 1.49, P < 10-5, N = 5 mice, 20 trials each. **(c)** DA timecourses after full-field optogenetic activation of CINs with a 0.5 sec pulse train across a variety of frequencies indicated at the top. Each trace is averaged across animals and displayed as Mean ± SEM. **(d)** Quantification of peak dopamine levels in early and late analysis windows for CIN stimulation frequency as indicated in Fig. 1c. The first DA peak is not significantly modulated by CIN stimulation frequency (one-way rmANOVA, main effect of stim frequency on early peak DA F(4,12) = 0.9, P = 0.47). By contrast, the second DA peak is significantly modulated by CIN stimulation frequency (one-way rmANOVA, main effect of stim frequency on late peak DA: F(4,12) = 4.8, P = 0.015). **(e)** Spectrogram of DA dynamics aligned to CIN stimulations, averaged across animals for baseline (top), DHβE administration (middle), and the subtraction of the two conditions. White traces are superimposed DA responses to facilitate comparing timecourses. Inset at bottom is an expansion of the first 100ms after laser onset, with a white arrow indicating power in frequency bands higher than 20-30Hz. **(f)** Normalized DA power spectra in the −0.5s to 2.5s window of optogenetic stimulation. The black line indicates the average for baseline trials without drug and blue line indicates trials with DHβE. Grey line is the subtraction of the two traces to highlight differences in specific frequency bands (shaded boxes, same color as in **g**) **(g)** Quantification of power at specific DA frequency bands. DHβE selectively attenuated power in beta and low gamma bands (two-way rmANOVA, significant main effect of drug: F(1,4) = 9.19, P = 0.038; drug x freqBand interaction: F(4,12) =10.96, P = 0.0001; post-hoc paired t-test: beta (t(4) = 3.92, P = 0.017) and low gamma (t(4) = 3.18, P = 0.033) bands are significantly different in DHBE than control. All other frequency bands have P > 0.15. **(h)** DA timecourses after full-field optogenetic activation of CINs with a 0.5 sec, 60Hz pulse train across a variety of spot sizes indicated at the top. Each trace is averaged across animals and displayed as Mean ± SEM. **(i)** Quantification of peak dopamine levels in early and late analysis windows for varying laser spot sizes as indicated in Fig. 1c. The first DA peak is significantly modulated by laser spot size one-way rmANOVA, main effect of spot size on first peak DA: F(4,12) = 16.8, P = 7 × 10^−5^). The second DA peak is also significantly increased by spot size: (one-way rmANOVA, main effect of stim frequency on late peak DA: F(4,12) = 8.25, P = 0.002). **(j)** Top: FOV of cannula illustrating quantification of DA fluorescence inside and outside of laser spot for an example animal. Bottom: Traces from three laser spot sizes showing the DA signals inside vs. outside the stimulated region. Red traces display the DA from inside the laser spot, gray traces show DA from outside the laser spot mask. **(k)** DA responses were detected outside the stimulated DS region. Peak DA magnitudes of the first and second peaks across spot sizes and inside vs. outside spot were not significantly different (2-way rmANOVA, effect of pixel location on β_2_-nAChR[DA] magnitude: F(1,12) = 4.38, P = 0.12).

We next assessed the temporal summation properties of CIN-evoked DA by stimulating CINs with trains of half-second laser pulses at varying frequencies. In contrast to the frequency-dependent magnitude scaling of the secondary DA peak, the magnitude and kinetics of ^β_2_-nAChR^[DA] peaks were insensitive to CIN stimulation frequency (**Fig. 4c-d**). This mirrors the all-or-none DHβE-sensitive DA response seen in slice CIN activation^9^ and indicates that ^β_2_-nAChR^[DA] retains conserved kinetics across diverse CIN firing regimes.

We reasoned that this remarkable shape invariance would generate a recognizable oscillatory footprint in the frequency domain of DA fluctuations. To examine this possibility, we performed time-frequency decomposition of the CIN-evoked DA signal and asked whether specific frequency bands display power modulation surrounding either DA peak, and if this band is attenuated with DHβE administration. Supporting this prediction, we observed a brief burst of power concentrated in the beta (15-25 Hz) and low gamma (30-60 Hz) frequency bands coincident with the arrival of the ^β_2_-nAChR^[DA] elevation (**Fig. 4e**). Notably, DHβE administration selectively diminished these bands, but not delta- or theta-band DA fluctuations (**Fig. 4f-g**). These results further bolster the presence of identifiable temporal signatures that uniquely index monosynaptic axonal regulation of DA via β_2_-nAChRs.

Finally, beyond these temporal features, ^β_2_-nAChR^[DA] showed a strong dependence on the spatial extent of synchronous CIN activation, as previously reported *ex vivo*^9^. When we restricted CIN stimulation to small DS quadrants, ^β_2_-nAChR^[DA] was absent, with only the second peak evoked (**Fig. 1e**). To quantify this spatial activation sensitivity, we systematically varied the laser spot diameter and found that ^β_2_-nAChR^[DA] required a minimum spatial scale of CIN synchrony and increased proportionally with larger stimulation fields (**Fig. 4h-i**;). Surprisingly, these DA responses were robustly detected outside the stimulated DS region, though mildly reduced (**Fig. 4j-k**), suggesting that once a spatial threshold of CIN recruitment is exceeded, ^β_2_-nAChR^[DA] is expressed as an all-or-none, DS-wide synchronized event. In contrast, the delayed DA elevation was robustly evoked even with the smallest stimulation size tested, though its amplitude was further increased with broader CIN recruitment (**Fig. 4h-i**). These results demonstrate that ^β_2_-nAChR^[DA] critically depends on the coordinated activation of DS CINs across a wide spatial domain, whereas the slower component can be initiated by localized CIN activity.

## Discussion

### Monosynaptic nicotinic control of dopamine is preserved in vivo

By combining high-speed fluorescence imaging with spatiotemporally precise optogenetic manipulations *in vivo*, we demonstrate that CIN activation elicits an ultra-fast DA elevation across the dorsal striatum that peaks within ~40 ms. The initial DA component exhibits all the defining hallmarks of monosynaptic axo-axonal cholinergic regulation of DA previously characterized *ex vivo*^7–10,42^: initiation within tens of milliseconds, strict dependence on β_2_-containing nicotinic receptors, and amplitude invariance across a broad range of CIN stimulation frequencies. These observations establish that the monosynaptic β_2_-nAChR-mediated regulation of DA axons is robustly expressed in awake behaving animals under conditions that permit large-scale CIN recruitment and high-bandwidth DA measurement.

### A polysynaptic CIN-DA interaction emerges in vivo to support the delayed peak

In parallel, CIN activation also evoked a second DA elevation with markedly slower kinetics, peaking ~180 ms after stimulation and persisting despite β_2_-nAChR blockade. This delayed component was reduced but not abolished by muscarinic receptor antagonism and was further attenuated by blockade of α_7_- and α_6_*-nAChRs, indicating recruitment via multiple redundant cholinergic pathways. Notably, a small but significant DA transient persisted even under simultaneous blockade of all nicotinic and muscarinic receptors, suggesting the presence of a non-cholinergic CIN messenger, likely glutamate^10,45,46^, that can promote DA release *in vivo*.

Importantly, this secondary DA transient is absent from slice studies, where DHβE eliminates all CIN-evoked DA release^8,9,43,44^, implying that this mechanism requires intact striatal-midbrain circuitry. Together, the properties of the second peak point to a parallel polysynaptic mechanism that is uniquely engaged *in vivo* and capable of supporting CIN-driven DA release even when the monosynaptic β_2_-nAChR mechanism is inaccessible or pharmacologically suppressed.

### Distinct spatiotemporal signatures reveal the basis of slice -vs- in vivo discrepancies

The two phases of CIN-evoked DA release exhibited distinct temporal, spectral, and spatial signatures. ^β_2_-nAChR^[DA] is marked by rapid onset, narrow duration, lack of temporal summation, and detected as a transient burst in the beta/low-gamma DA spectrum. By contrast, the delayed DA peak shows slower kinetics and graded dependence on CIN recruitment strength and spatial extent. These differences likely arise because synchronous CIN activation produces near-simultaneous excitation of most DA axons, yielding a spatially widespread axonal spike and DA efflux that is curtailed by rapid β_2_-nAChR desensitization and a short refractory period in DA axons. Together, these events will restrict the temporal width of ^β_2_-nAChR^[DA], preventing DA release from summating repeated CIN activations or incoming midbrain spikes propagat through DA axons, thereby appearing as a frequency filter on DA release^9,55^.

Spatially, ^β_2_-nAChR^[DA] required large-scale CIN synchrony, whereas the delayed peak could be triggered by stimulating CINs in small DS territories, suggesting that β_2_-nAChRs on DA axons read out the temporal coincidence of cholinergic drive rather than overall cholinergic tone, which may be achieved by other cholinergic receptors on DA axons^56,57^. The separable spatiotemporal signatures we identify here provide a principled framework for identifying ^β_2_-nAChR^[DA] events outside of optogenetic stimulation. They also clarify why previous *in vivo* studies did not detect β_2_-nAChR-dependent DA modulation^37,39,40^: We show that either limited temporal bandwidth or slow sensor kinetics alone collapses the two CIN-evoked components into a single sluggish transient, occluding the defining temporal structure and pharmacological properties of ^β_2_-nAChR^[DA]. Together with the stringent spatial scale required for synchronous CIN recruitment and the need to mitigate optical artifacts^38^, these factors and properties of ^β_2_-nAChR^[DA] dynamics account for the apparent discrepancy between slice studies and recent *in vivo* assessments of β_2_-nAChR-dependent DA release.

### Dual cholinergic pathways expand the computational repertoire of striatal DA

Our identification of two mechanistically separable CIN-evoked DA signals *in vivo* bolsters longstanding evidence that CINs directly promote striatal DA release, while challenging the view that this interaction is dominated by a single nicotinic, axo-axonal mechanism. Historically, slice preparations predominantly revealed monosynaptic CIN-evoked DA changes which were completely blocked by DHβE^7–10,43,44^ (though other direct mechanisms are also reported^56–58^), prompting previous hypotheses to focus exclusively on this mechanism when considering computations supported by cholinergic regulation of DA^8,30,31,37,39,40,55,59–61^. Our findings reveal that while the ultra-fast β_2_-nAChR mechanism is indeed expressed *in vivo*, it is only one component of CIN-evoked DA release. The delayed polysynaptic DA elevation we report here has not been described previously *in vivo* and is inaccessible to *ex vivo* characterization. Yet, it provides a parallel, temporally extended, and spatially flexible mode of cholinergic control that closely aligns with the hypothesized behavioral contributions of DA to online motivational and cognitive vigor^30–32,36,37^.

We posit that the dual architecture we characterized reframes CINs as a multi-channel DA regulator capable of driving both rapid, global circuit resets and spatially localized modulation of DA across performance epochs. Specifically, CIN-mediated nicotinic activation produces a spatiotemporally all-or-none global DA event consistent with a circuit-wide ‘interrupt’ signal capable of rapidly resetting striatal computations on millisecond timescales. Because this ^β_2_-nAChR^[DA] mode is temporally compressed, frequency invariant, and requires synchronous CIN recruitment across a broad anatomical domain, it is likely engaged only during rare, sharply disruptive behavioral events. Such a mechanism is ideally positioned to prompt wholesale policy shifts when latent task rules change^62–64^, or to swiftly reorient ongoing motor programs^65^, for example, toward escaping aversive events^66^. In contrast, the delayed DA elevation reflects a qualitatively different regime of CIN regulation of DA dynamics that is slower, spatially graded, receptor-redundant, and robust to pharmacological blockade of β_2_-nAChRs. These properties are incompatible with direct axonal activation and instead point toward a polysynaptic relay embedded within the striatum-midbrain loop. A plausible circuit is a concerted effort by CINs to activate D1-expressing medium spiny neurons^45,49,67^, which in turn disinhibits midbrain DA neurons by inhibiting the SNr^68^. This architecture would allow CINs to modulate DA release indirectly, even when nicotinic mechanisms are desensitized or saturated, and would generate DA that is ultimately midbrain-sourced, but locally specified by striatal interactions. The delayed transient’s slower kinetics, tolerance to extensive cholinergic blockade, and recruitment by small CIN territories position it as an ideal mechanism for CIN-evoked DA modulation of flexible approach behaviors^26,69,70^, local ensemble competition^71–73^, or sustaining performance-epoch DA ramps without desensitization^25–27^. Thus, the second peak represents a previously unrecognized, polysynaptic route by which CINs influence DA on behaviorally relevant timescales.

In summary, the β_2_-nAChR and delayed polysynaptic components reveal that CINs regulate striatal dopamine through two pharmacologically distinct mechanisms. β_2_-nAChRs provide a monosynaptic, temporally precise, and spatially expansive mechanism for rapid DA efflux, whereas the delayed polysynaptic mode supplies a slower, spatially flexible pathway for shaping DA dynamics during ongoing behavior. These complementary channels expand the repertoire of DA computations, supporting an architecture in which CINs and midbrain DA neurons jointly coordinate behavioral control through dual cholinergically initiated DA pathways operating on separate timescales. Our studies offer a mechanistically grounded framework for modular RL algorithms that can be mobilized under specific behavioral demands.

## Acknowledgements

We thank Janet Dubinsky for feedback on earlier versions of the manuscript and members of the Hamid Lab for feedback at various stages of the project. We also thank Julia Lemos for sharing pharmacological resources. This work was supported by awards from the Howard Hughes Medical Institute (Hanna Gray Fellowship to AAH), the National Institute of Health (P50MH119569-06 to AAH; P30DA048742 to AAH; T32GM156735-01 and T32DA007234 to NGF), UMN Dean’s office (ECRA award to AAH), and UMN Medical Discovery Team on Addiction.

## Author Contributions

DRF collected the data and contributed to data analysis and writing. NGF and HZ assisted with data analysis. AAH designed and supervised the study, analyzed the data, and wrote the manuscript.

## Data and code availability

All data and code are available from the corresponding author(s) upon reasonable request.

## Materials and Correspondence

Correspondence and material requests are to be addressed to Arif Hamid.

## Methods

### Animals and Surgery

Adult male and female ChAT-cre+ mice(JAX, 006410) were group housed on a reversed 12-hour cycle, and all experiments were performed during the dark phase. All procedures were approved by the University of Minnesota Institutional Animal Care and Use Committee and conformed to NIH guidelines.

All stereotaxic surgeries were performed under aseptic conditions. Mice were anesthetized with isoflurane (5% for induction, 1-2% for maintenance in 1 L/min oxygen), and body temperature was maintained at 37°C with a heating pad. To achieve cell-type-specific expression of red-shifted opsin and fluorescent green DA sensors in the striatum, ChAT-Cre^+^ mice received a stereotactic viral injection of cre-dependent ChrimsonR (pAAV-Syn-FLEX-rc[ChrimsonR-tdTomato] AAV5, addgene cat#62723) together with dLight1.2 (pAAV-hsyn-dlight1.2 AAV5, addgene cat#111068) or GRAB-DA2m (pAAV-hsyn_GRAB_DA2m AAV9, addgene cat#140553) into the dorsal striatum. Viral injections were delivered through burr holes targeting the DS at stereotaxic coordinates (in mm): AP 0.7, 0.7, −0.5, ML 1.5, 2.5, 2, and 0.5 µL of virus was injected at DV −2.6. This injection scheme yielded widespread dorsal striatal expression of both the cre-dependent ChrimsonR and dLight1.2 or GRAB-DA2m constructs. Immediately after viral injection, a custom stainless-steel headpost was cemented to the mouse’s skull.

After a 3-week viral expression period, mice underwent a second surgery to implant an imaging cannula at the site of DS injections as described previously^36^. Briefly, a 3-mm craniotomy was made over the DS, followed by careful resection of the dura and overlying cortex. Cortical tissue above the dorsal striatum was slowly aspirated to minimize bleeding and cellular damage and optimize optical access. A custom cannula was constructed by bonding a circular coverslip to a 3-mm diameter, 2.5 mm tall stainless-steel cylinder using Norland Optical Adhesive. This cannula was then positioned so that the coverslip rested directly on the exposed dorsal striatum and was secured to the skull with dental cement. Following all surgeries, mice received a 3-day post-operative care regimen consisting of daily health checks, subcutaneous analgesia, and weight monitoring. At the end of the experiments, mice were perfused, and brains were processed for histological analysis to verify viral expression and cannula placement.

### Widefield Imaging, optogenetic stimulation, and systemic pharmacology

Fluorescence imaging was performed on a custom-built wide-field imaging system that supported simultaneous optical stimulation and imaging through the implanted cannula^36^. Briefly, dLight fluorescence was excited using continuous 470-nm LED illumination. This excitation light was reflected by a 425SP dichroic mirror, transmitted through a 490SP long-pass dichroic mirror, and directed through a 4X Thorlabs objective onto the dorsal striatum. Emitted green fluorescence returned to the camera via a 525±50nm bandpass emission filter before digitization by either a PCO Edge 5.5 camera or a FLIR Grasshopper (GS3-U3-32S4M) camera. To deliver spatiotemporally specific optogenetic stimulation to the dorsal striatum, a fiber-coupled 635nm laser (LSR-040-0463-0635, Mightex) was triggered by custom LABVIEW control software and routed through a digital micromirror device (DMD; Polygon1000-DL, Mightex). The DMD received independent patterns that defined the spatial mask of laser stimulation. After reflection from the DMD, this patterned 635nm light was combined with the excitation path via the 490nm long-pass dichroic and projected on the striatum through the objective. An emission filter in the detection path prevented this laser light from reaching the camera. We regularly calibrated the stimulation system to maintain alignment between the DMD projection and the cannula’s field of view, as well as the spatial distribution and power density at each projected pixel. All experiments described used an analog voltage drive of the laser to achieve 50-55mW at the objective, which translated to 7.5 mW/mm^2^ across the 7mm^2^ FOV of the imaging cannula when all the pixels of the DMD were set to reflection. Spot size manipulation experiments described in Fig. 1g-h and Fig. 4h manipulated the fraction of DMD mirrors reflecting laser light at the same TTL voltage, thus yielding the same 7.5 mW/mm^2^, irrespective of the spot size projected.

We used custom LabVIEW scripts to systematically vary the parameters of excitation LEDs, laser activation, DMD projection, and camera exposure, and to store various behavioral and metadata associated with each experiment. A subset of the data was collected under continuous excitation LED and frame capture throughout the session with optogenetic stimulations randomly delivered 10-30 seconds apart, either full-field, at quadrants of DS, or concentric laser spots, depending on experimental condition. Session with dLight1.2 and GRAB.DA2m imaging at 500Hz employed a triggered excitation LED approach to minimize signal degradation from continuous fluorophore excitation. On each trial, the excitation LED is illuminated for a random 5-15-second interval before capturing frames for a three-second duration, where the laser is activated after a half-second baseline period. The laser was activated by a stimulus train designed piecemeal in LABVIEW for a TTL of a specified number of pulses, duration, and frequency. Each session lasted 60-90 minutes to capture a minimum of 20 trials for each condition.

All pharmacological experiments were performed using intraperitoneal (i.p.) injections, either 10 minutes before imaging began or at the halfway point. Pharmacological experiments that examined the combined effects of multiple receptor types were premixed in a container prior to i.p. administration. The respective receptor contributions to cholinergic evoked DA release used the following antagonists: Dihydro-β-erythroidine hydrobromide (DHβE) for β_2_-subunit-containing nicotinic receptor blockade (1 mg/kg; Tocris cat#: 234910); scopolamine for broad muscarinic receptor blockade (1 or 5 mg/kg; Sigma-Aldrich cat#: S0929); mecamylamine for broad nicotinic receptor blockade blockade (1 or 5 mg/kg; Sigma-Aldrich cat#: M9020); Methyllycaconitine citrate (MLA) for α_7_-nicotinic receptor blockade (1mg/kg; MedChem Express cat#: HY-N2332A); 1,1’-(decane-1,10-diyl) bis (3-methylpyridin-1-ium) iodide (bPiDI) for α_6_*-nicotinic receptor blockade (1mg/kg; MedChem Express cat#: HY-152170).

### Data Analysis and statistical comparisons

Widefield dLight imaging data were initially stored as 8- or 16-bit TIFF stacks and processed in MATLAB using custom routines. Processing steps included application of a striatal mask, spatial registration, temporal synchronization with stimulation or behavioral events, and conversion into high-dimensional HDF5 files for downstream analyses. Unless otherwise specified, fluorescence responses were averaged across all striatal pixels to yield a single dorsal striatum (DS) time series for each trial; in some analyses (e.g., Fig. 1f–h, 3b, 4j–k), pixelwise or regional averages were computed using masks defined by the optogenetic stimulation masks.

To compare CIN-evoked dopamine responses across trials, mice, and pharmacological conditions, fluorescence traces were normalized within trial to a pre-stimulation baseline. For each trial, the mean and standard deviation of the baseline window were computed, and all subsequent frames were expressed as z-scores. Latency to fluorescence onset (Fig. 1e) was defined as the first post-stimulus time point at which the z-scored trace exceeded one baseline standard deviation. Latency to peak was measured using convergent algorithms, including derivative-based curvature assessment and MATLAB’s *findpeaks* function applied within predefined early and late peak windows. Area-under-the-curve (AUC) measurements were computed by integrating the z-scored signal within the corresponding temporal windows. The rate of rise (Fig. 3d) was defined as the average change in z-score per second during the rising phase.

To characterize the sharpness of the rising phase of early and late CIN-evoked dopamine peaks, we estimated an exponential rise time constant τ. For each trial, baseline-subtracted and z-scored traces were truncated from stimulus onset to the detected peak, and only positive-valued samples were retained to isolate the monotonic rise. Assuming an exponential form DA(t) = A·exp(t/τ), we linearized the model via natural logarithm and fit log(DA) with linear least squares; τ was computed as the inverse of the fitted slope. FWHM was computed for each dopamine transient by identifying the half-amplitude value defined as the midpoint between baseline fluorescence and peak amplitude. The rising and falling time points at which the trace first crossed, and later returned to, this half-amplitude threshold were identified. FWHM was measured as the temporal difference between these two crossings.

Spectral decomposition of DA fluorescence signals was performed using Welch’s method (*pwelch* function in MATLAB), yielding power density estimates from 0 to 250 Hz based on a 500 Hz sampling rate. Time-frequency representations were generated using a custom short-time Fourier decomposition, and spectrograms were averaged across trials and across mice to visualize stimulus-locked spectral dynamics (Fig. 4e-f).To quantify power within specific physiologically relevant frequency bands, we integrated the Welch spectrum over predefined frequency bands (0.5-4 Hz for Delta band, 6-10 Hz for Theta band,15-30 Hz for Beta band, 30-60 Hz for Low-Gamma band,70-90 Hz for High-Gamma band) for each mouse and compared baseline versus DHβE conditions within-mouse (Fig. 4g).

For statistical tests, we used MATLAB implementation of non-parametric methods that do not assume underlying distributions of data, or analyses of variance paired with simple post-hoc ttests.

## References

1. Liu, C., Goel, P. & Kaeser, P. S. Spatial and temporal scales of dopamine transmission. Nat. Rev. Neurosci. 22, 345–358 (2021).

2. Maia, T. V. & Frank, M. J. An Integrative Perspective on the Role of Dopamine in Schizophrenia. Biol. Psychiatry 81, 52–66 (2017).

3. Gerfen, C. R. & Surmeier, D. J. Modulation of striatal projection systems by dopamine. Annu. Rev. Neurosci. 34, 441–466 (2011).

4. Todd, K. L., Cramb, K. M. L., Brimblecombe, K. R. & Cragg, S. J. New insights into axonal regulators of dopamine transmission in health and disease. Curr. Opin. Neurobiol. 94, 103093 (2025).

5. Cragg, S. J., Sulzer, D. & Todd, K. Axonal regulation of dopamine transmission by striatal neuromodulators. *Handb*. Behav. Neurosci. (2025).

6. Holly, E. N., Galanaugh, J. & Fuccillo, M. V. Local regulation of striatal dopamine: A diversity of circuit mechanisms for a diversity of behavioral functions? 85, 102839 (2024).

7. Kramer, P. F. et al. Synaptic-like axo-axonal transmission from striatal cholinergic interneurons onto dopaminergic fibers. Neuron 110, 2949–2960.e4 (2022).

8. Liu, C. et al. An action potential initiation mechanism in distal axons for the control of dopamine release. Science 375, 1378–1385 (2022).

9. Threlfell, S. et al. Striatal dopamine release is triggered by synchronized activity in cholinergic interneurons. Neuron 75, 58–64 (2012).

10. Cachope, R. et al. Selective activation of cholinergic interneurons enhances accumbal phasic dopamine release: setting the tone for reward processing. Cell Rep. 2, 33–41 (2012).

11. Kosillo, P., Zhang, Y.-F., Threlfell, S. & Cragg, S. J. Cortical Control of Striatal Dopamine Transmission via Striatal Cholinergic Interneurons. Cereb. Cortex 26, 4160–4169 (2016).

12. Ding, J. B., Guzman, J. N., Peterson, J. D., Goldberg, J. A. & Surmeier, D. J. Thalamic gating of corticostriatal signaling by cholinergic interneurons. Neuron 67, 294–307 (2010).

13. Mandelbaum, G. et al. Distinct Cortical-Thalamic-Striatal Circuits through the Parafascicular Nucleus. Neuron 102, 636–652.e7 (2019).

14. Adrover, M. F. et al. Prefrontal cortex driven dopamine signals in the striatum show unique spatial and pharmacological properties. J. Neurosci. (2020) doi:10.1523/JNEUROSCI.1327-20.2020.

15. Guo, Q. et al. Whole-brain mapping of inputs to projection neurons and cholinergic interneurons in the dorsal striatum. PLoS One 10, e0123381 (2015).

16. Schultz, W., Dayan, P. & Montague, P. R. A neural substrate of prediction and reward. Science 275, 1593–1599 (1997).

17. Steinberg, E. E. et al. A causal link between prediction errors, dopamine neurons and learning. Nat. Neurosci. 16, 966–973 (2013).

18. Eshel, N., Tian, J., Bukwich, M. & Uchida, N. Dopamine neurons share common response function for reward prediction error. Nat. Neurosci. 19, 479–486 (2016).

19. Matsuda, W. et al. Single nigrostriatal dopaminergic neurons form widely spread and highly dense axonal arborizations in the neostriatum. J. Neurosci. 29, 444–453 (2009).

20. Aransay, A., Rodríguez-López, C., García-Amado, M., Clascá, F. & Prensa, L. Long-range projection neurons of the mouse ventral tegmental area: a single-cell axon tracing analysis. Front. Neuroanat. 9, 59 (2015).

21. Schultz, W. Predictive reward signal of dopamine neurons. J. Neurophysiol. 80, 1–27 (1998).

22. Glimcher, P. W. Understanding dopamine and reinforcement learning: the dopamine reward prediction error hypothesis. Proc. Natl. Acad. Sci. U. S. A. 108 Suppl 3, 15647–15654 (2011).

23. Kim, H. R. et al. A Unified Framework for Dopamine Signals across Timescales. Cell (2020) doi:10.1016/j.cell.2020.11.013.

24. Gershman, S. J. et al. Explaining dopamine through prediction errors and beyond. Nat. Neurosci. 27, 1645–1655 (2024).

25. Collins, A. L. et al. Dynamic mesolimbic dopamine signaling during action sequence learning and expectation violation. Sci. Rep. 6, 20231 (2016).

26. Hamid, A. A. et al. Mesolimbic dopamine signals the value of work. Nat. Neurosci. 19, 117–126 (2016).

27. Mohebi, A. et al. Dissociable dopamine dynamics for learning and motivation. Nature 570, 65–70 (2019).

28. Westbrook, A. et al. Dopamine promotes cognitive effort by biasing the benefits versus costs of cognitive work. Science 367, 1362–1366 (2020).

29. Howe, M. W., Tierney, P. L., Sandberg, S. G., Phillips, P. E. M. & Graybiel, A. M. Prolonged dopamine signalling in striatum signals proximity and value of distant rewards. Nature 500, 575–579 (2013).

30. Berke, J. D. What does dopamine mean? Nat. Neurosci. 21, 787–793 (2018).

31. Hamid, A. A. Dopaminergic specializations for flexible behavioral control: linking levels of analysis and functional architectures. Current Opinion in Behavioral Sciences 41, 175–184 (2021).

32. Frank, M. J. Adaptive cost-benefit control fueled by striatal dopamine. Annu. Rev. Neurosci. 48, (2025).

33. Gottschalk, A. C., Faturos, N. G. & Hamid, A. A. Timescales of dopamine release in the striatum as a window into hierarchical control. 66, 101589 (2025).

34. Franklin, N. T. & Frank, M. J. A cholinergic feedback circuit to regulate striatal population uncertainty and optimize reinforcement learning. Elife 4, (2015).

35. Kim, T. et al. The functional role of striatal cholinergic interneurons in reinforcement learning from computational perspective. Front. Neural Circuits 13, 10 (2019).

36. Hamid, A. A., Frank, M. J. & Moore, C. I. Wave-like dopamine dynamics as a mechanism for spatiotemporal credit assignment. Cell (2021) doi:10.1016/j.cell.2021.03.046.

37. Mohebi, A., Collins, V. L. & Berke, J. D. Accumbens cholinergic interneurons dynamically promote dopamine release and enable motivation. Elife 12, (2023).

38. Taniguchi, J. et al. Comment on ‘Accumbens cholinergic interneurons dynamically promote dopamine release and enable motivation’. eLife 13, e95694 (2024).

39. Krok, A. C. et al. Intrinsic dopamine and acetylcholine dynamics in the striatum of mice. Nature 621, 543–549 (2023).

40. Chantranupong, L. et al. Dopamine and glutamate regulate striatal acetylcholine in decision-making. Nature 621, 577–585 (2023).

41. Kuan, L. et al. Neuroinformatics of the Allen Mouse Brain Connectivity Atlas. Methods 73, 4–17 (2015).

42. Wang, L. et al. Temporal components of cholinergic terminal to dopaminergic terminal transmission in dorsal striatum slices of mice: Timing of axo-axonal striatal dopaminergic transmission. J. Physiol. 592, 3559–3576 (2014).

43. Mamaligas, A. A., Cai, Y. & Ford, C. P. Nicotinic and opioid receptor regulation of striatal dopamine D2-receptor mediated transmission. Sci. Rep. 6, 37834 (2016).

44. Brimblecombe, K. R. et al. Targeted activation of cholinergic interneurons accounts for the modulation of dopamine by striatal nicotinic receptors. eNeuro 5, ENEURO.0397–17.2018 (2018).

45. Higley, M. J. et al. Cholinergic interneurons mediate fast VGluT3-dependent glutamatergic transmission in the striatum. PLoS One 6, e19155 (2011).

46. Nelson, A. B., Bussert, T. G., Kreitzer, A. C. & Seal, R. P. Striatal cholinergic neurotransmission requires VGLUT3. J. Neurosci. 34, 8772–8777 (2014).

47. Gras, C. et al. The vesicular glutamate transporter VGLUT3 synergizes striatal acetylcholine tone. Nat. Neurosci. 11, 292–300 (2008).

48. Assous, M. Striatal cholinergic transmission. Focus on nicotinic receptors’ influence in striatal circuits. Eur. J. Neurosci. 53, 2421–2442 (2021).

49. Campos, F., Alfonso, M. & Durán, R. In vivo modulation of alpha7 nicotinic receptors on striatal glutamate release induced by anatoxin-A. Neurochem. Int. 56, 850–855 (2010).

50. Howe, W. M., Young, D. A., Bekheet, G. & Kozak, R. Nicotinic receptor subtypes differentially modulate glutamate release in the dorsal medial striatum. Neurochem. Int. 100, 30–34 (2016).

51. Tanimura, A., Du, Y., Kondapalli, J., Wokosin, D. L. & Surmeier, D. J. Cholinergic interneurons amplify thalamostriatal excitation of striatal indirect pathway neurons in Parkinson’s disease models. Neuron 101, 444–458.e6 (2019).

52. Labouesse, M. A., Cola, R. B. & Patriarchi, T. GPCR-based dopamine sensors-A detailed guide to inform sensor choice for in vivo imaging. Int. J. Mol. Sci. 21, 8048 (2020).

53. Patriarchi, T. et al. Ultrafast neuronal imaging of dopamine dynamics with designed genetically encoded sensors. Science 360, (2018).

54. Sun, F. et al. Next-generation GRAB sensors for monitoring dopaminergic activity in vivo. Nat. Methods 17, 1156–1166 (2020).

55. Zhang, Y.-F. et al. An axonal brake on striatal dopamine output by cholinergic interneurons. Nature Neuroscience 28, 783–794 (2024).

56. Shin, J. H., Adrover, M. F., Wess, J. & Alvarez, V. A. Muscarinic regulation of dopamine and glutamate transmission in the nucleus accumbens. Proc. Natl. Acad. Sci. U. S. A. 112, 8124–8129 (2015).

57. Razidlo, J. A. et al. Chronic loss of muscarinic M5 receptor function manifests disparate impairments in exploratory behavior in male and female mice despite common dopamine regulation. J. Neurosci. 42, 6917–6930 (2022).

58. Threlfell, S. et al. Striatal muscarinic receptors promote activity dependence of dopamine transmission via distinct receptor subtypes on cholinergic interneurons in ventral versus dorsal striatum. Journal of Neuroscience 30, 3398–3408 (2010).

59. Cragg, S. J. Meaningful silences: how dopamine listens to the ACh pause. Trends Neurosci. 29, 125–131 (2006).

60. Cachope, R. & Cheer, J. F. Local control of striatal dopamine release. Front. Behav. Neurosci. 8, 188 (2014).

61. Touponse, G. C. et al. Cholinergic modulation of dopamine release drives effortful behavior. bioRxivorg 2025.06.18.660394 (2025) doi:10.1101/2025.06.18.660394.

62. Nassar, M. R., Wilson, R. C., Heasly, B. & Gold, J. I. An approximately Bayesian delta-rule model explains the dynamics of belief updating in a changing environment. J. Neurosci. 30, 12366–12378 (2010).

63. Mah, A., Golden, C. E. M. & Constantinople, C. M. Dopamine transients encode reward prediction errors independent of learning rates. Cell Rep. 43, 114840 (2024).

64. Soltani, A. & Izquierdo, A. Adaptive learning under expected and unexpected uncertainty. Nat. Rev. Neurosci. 20, 635–644 (2019).

65. Howe, M. et al. Coordination of rapid cholinergic and dopaminergic signaling in striatum during spontaneous movement. Elife 8, (2019).

66. Macdonald, E. E. et al. A synaptic mechanism for encoding the learned value of action-derived safety. bioRxivorg 2025.08.04.668532 (2025) doi:10.1101/2025.08.04.668532.

67. Shen, W., Hamilton, S. E., Nathanson, N. M. & Surmeier, D. J. Cholinergic suppression of KCNQ channel currents enhances excitability of striatal medium spiny neurons. J. Neurosci. 25, 7449–7458 (2005).

68. Ambrosi, P. & Lerner, T. N. Striatonigrostriatal circuit architecture for disinhibition of dopamine signaling. Cell Rep. 40, 111228 (2022).

69. Nicola, S. M. The Flexible Approach Hypothesis: Unification of Effort and Cue-Responding Hypotheses for the Role of Nucleus Accumbens Dopamine in the Activation of Reward-Seeking Behavior. J. Neurosci. 30, 16585–16600 (2010).

70. du Hoffmann, J. & Nicola, S. M. Dopamine invigorates reward seeking by promoting cue-evoked excitation in the nucleus accumbens. J. Neurosci. 34, 14349–14364 (2014).

71. Dobbs, L. K. et al. Dopamine regulation of lateral inhibition between striatal neurons gates the stimulant actions of cocaine. Neuron 90, 1100–1113 (2016).

72. Tecuapetla, F., Koós, T., Tepper, J. M., Kabbani, N. & Yeckel, M. F. Differential dopaminergic modulation of neostriatal synaptic connections of striatopallidal axon collaterals. J. Neurosci. 29, 8977–8990 (2009).

73. Carrillo-Reid, L., Hernández-López, S., Tapia, D., Galarraga, E. & Bargas, J. Dopaminergic modulation of the striatal microcircuit: receptor-specific configuration of cell assemblies. J. Neurosci. 31, 14972–14983 (2011).

